# Transcriptomes and Raman spectra are linked linearly through a shared low-dimensional subspace

**DOI:** 10.1101/235580

**Authors:** Koseki J. Kobayashi-Kirschvink, Hidenori Nakaoka, Arisa Oda, Ken-ichiro F. Kamei, Kazuki Nosho, Hiroko Fukushima, Yu Kanesaki, Shunsuke Yajima, Haruhiko Masaki, Kunihiro Ohta, Yuichi Wakamoto

**Author notes:** Correspondence to: koseki or Lead contact: Y.W.

## Abstract

Raman spectroscopy is an imaging technique that can reflect whole-cell molecular compositions *in vivo,* and has been applied recently in cell biology to characterize different cell types and states. However, due to the complex molecular compositions and spectral overlaps, the interpretation of cellular Raman spectra have remained unclear. In this report, we compared cellular Raman spectra to transcriptomes of *Schizosaccharomyces pombe* and *Escherichia coli,* and provide firm evidence that they can be computationally connected and interpreted. Specifically, we find that the dimensions of high-dimensional Raman spectra and transcriptomes measured by RNA-seq can be effectively reduced and connected linearly through a shared low-dimensional subspace. Accordingly, we were able to reconstruct global gene expression profiles by applying the calculated transformation matrix to Raman spectra, and vice versa. Strikingly, highly expressed ncRNAs contributed to the Raman-transcriptome linear correspondence more significantly than mRNAs in *S. pombe,* which implies their major role in coordinating molecular compositions. This compatibility between whole-cell Raman spectra and transcriptomes marks an important and promising step towards establishing spectroscopic live-cell omics studies.

## Introduction

Raman spectroscopy is a laser-based analytical technique that measures the energy shift of scattered photons caused by molecular bond vibrations. Specific molecules have unique Raman spectral signatures, which in turn allows us to determine the chemical species in target samples. This technique is applicable to biological samples, and can potentially unravel the abundances of various biomolecules in cells and tissues in a comprehensive, non-destructive, and label-free manner.

Typically, interpreting spectra involves decomposing them into those of known purified spectra and quantifying the corresponding molecules. Numerous methods such as multivariate curve resolution alternating least squares (MCR-ALS) have been developed [1, 2]. However, preparing spectra of each and every biomolecule of cells is laborious, or even impossible. In addition, none of these methods resolve severe spectral overlaps of biomolecules, which makes unique quantification intractable. Consequently, it is widely recognized that discerning constituent molecular species in a comprehensive manner is difficult, making the interpretation of whole-cell Raman spectra nearly intractable [3–6].

Alternative approaches of interpretation are to represent intrinsically high-dimensional cellular Raman spectra in low-dimensional spaces using dimension reduction methods such as principal component analysis (PCA) and linear discriminant analysis (LDA) [1, 7, 8]. Though these methods can sometimes successfully assign the spectra from different cell types or states to distinct subspaces, the interpretation still remains unclear because the resulting axes and spaces usually cannot be characterized by any biological properties. These approaches therefore often fail to provide any mechanistic insights into the differences of the spectra.

In this report, instead of pursuing the spectral decomposition, we asked whether whole-cell Raman spectra could be directly and computationally corresponded to other types of well studied omics-level information. Employing dimension reduction methods, we reveal a surprising correspondence between cellular Raman spectra and transcriptomes for *S. pombe* and *E. coli*. We show that a simple linear transformation links these two types of high-dimensional data, and demonstrate that global expression profiles of transcriptomes across culture conditions can be reconstructed in non-destructive manners from cellular Raman spectra, which was made possible by the intrinsic low-dimensionality of transcriptomes. Furthermore, interestingly in *S. pombe,* ncRNAs contributed to the Raman-transcriptome linearity more significantly than mRNAs, supporting their major role in coordinating total molecular compositions in eukaryotic cells. Together, these results show that whole-cell Raman spectra can be directly and computationally linked to cellular omics information, and paves a new way to conducting spectroscopic live-cell omics studies in the future.

## Results

### PC-LDA can reveal distinct cellular states of *S. pombe* from Raman spectra

We obtained Raman spectra of single *S. pombe* cells sampled from 10 different culture conditions using a custom-built Raman microscope with 532 nm excitation wavelength, 10 s exposure time and 4 mW power at the sample stage (Fig. S1). Technical details on signal filtering and noise reduction are explained in Materials and Methods. Culture conditions are listed in Table S1, which includes rich and minimal media, nutrient depleted media, and various stress conditions. Prior to measurements of Raman spectra, cells were fixed with 2% formaldehyde at 4°C. We obtained Raman spectra from 54-76 cells per condition.

Raman spectra from cells had common features: the strong signal peaks of CH_2_ and CH_3_ bonds around 2800 to 3000 cm^−1^; the silent region from 1800 cm^−1^ to 2800 cm^−1^; and the rugged peaks from 700 cm^−1^ to 1800 cm^−1^ (Fig. 1A, IB, and S2). These global features are common among various cell types including mammalian cells [3,9–11], thus reflecting the basic chemical composition of cells. The spectral range from 700 cm^−1^ to 1800 cm^−1^, the *fingerprint region of biological samples* [9, 10], is where most of the signals such as proteins and metabolites are observed. We therefore focused on this spectral region in the following analyses.

**Figure 1.**
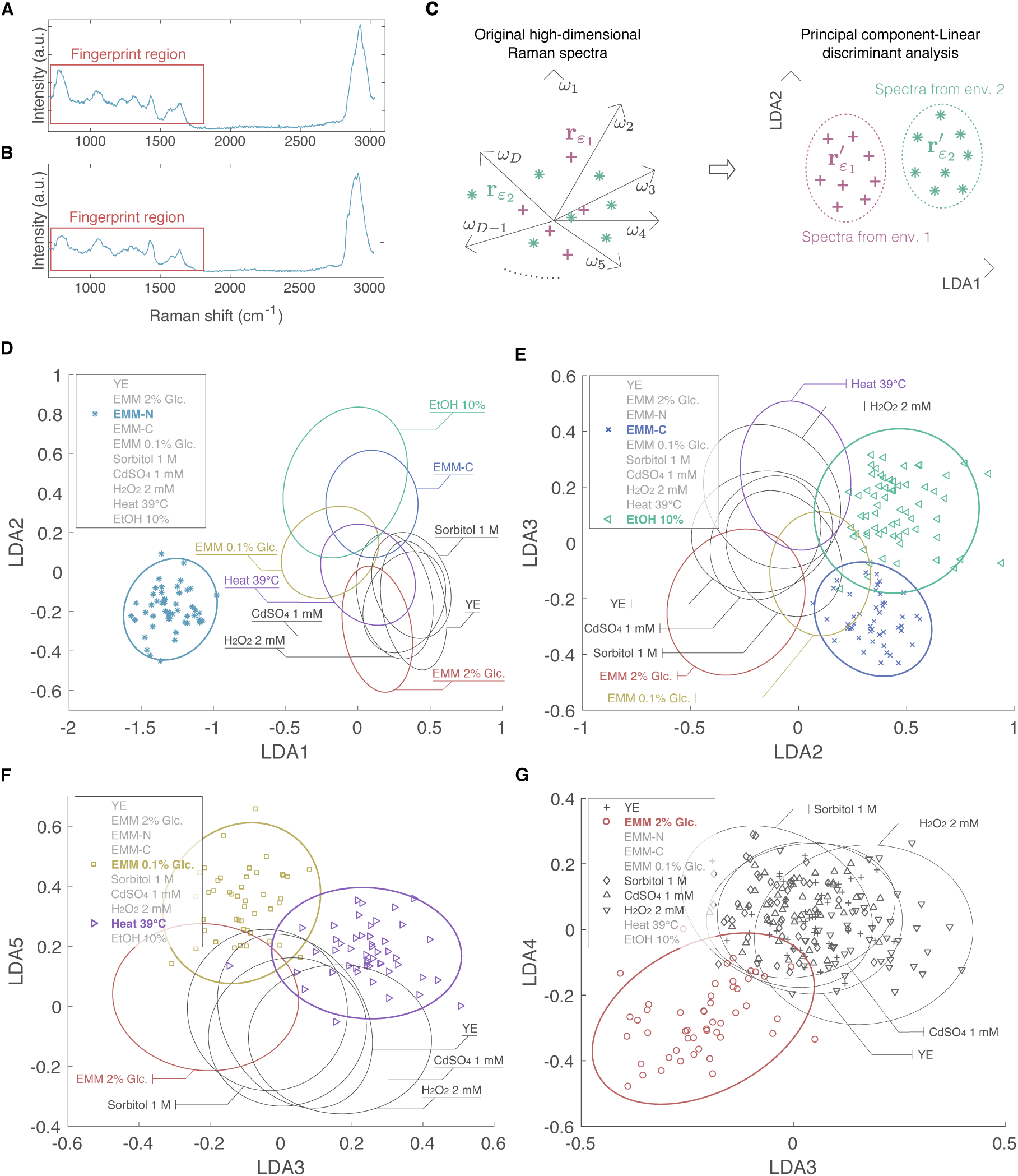
Measurement and dimension reduction of single-cell Raman spectra of *S. pombe*. A, B. Raman spectra of single cells cultured in rich medium (YE, plot A) and nitrogen-depleted medium (EMM-N, plot B). The fingerprint region of the spectra from 700 cm^−1^ to 1800 cm^−1^ is indicated by red rectangles. C. Dimension reduction of Raman spectra. Raman spectrum from a single cell in environment *ε* can be expressed as a single point *r_ε_* in a high-dimensional space whose axes *ω_i_* represent the signal intensity at specific Raman shift positions. Principal component-linear discriminant analysis (PC-LDA) is applied to Raman spectra to remove systematic-error while simultaneously reducing the dimensionality. If environmentally-dependent spectral features exist, PC-LDA can assign spectra from different environments to unique different clusters in low-dimensional LDA spaces. **D-G.** Single-cell Raman spectra processed by PC-LDA and expressed in low-dimensional space (ellipses, the *χ^2^* 95% confidence intervals of the mean of each condition). Notably, the first few LDA axes were able to distinguish spectra from cells cultured in nitrogen-depleted medium (EMM-N), ethanol stress (EtOH 10%), glucose depleted (EMM-C), glucose limited (EMM 0.1% Glc.), heat shock (Heat 39°C) and glucose-supplemented minimal medium (EMM 2% Glc.).

We first asked whether these spectra can be classified based on the culture condition from which the cells were sampled, and conducted the principal component-linear discriminant analysis (PC-LDA) [7, 8, 12]. A Raman spectrum from a cell can be represented as a single point in a high-dimensional space (599 dimensions in our measurements) where the signal intensity at a specific Raman-shift wavenumber position corresponds to one dimension (Fig. 1C). Taking the culture-condition assignments into account and simultaneously avoiding over-fitting, PC-LDA computationally extracts the most discriminatory bases by maximizing the ratio of the between-group variances to the sum of within-group variances in the lower dimensional representation (Fig. 1C, and see Materials and Methods). PC-LDA reduces the dimensions to the number of groups (environments) –1; we therefore reduced the dimensions of Raman spectra to 9 in our analysis.

Our results show that Raman spectra from the same condition form clusters in the dimension-reduced Raman space (Fig. 1D-G). Some of the clusters could be recognized by the first few LDA axes. Most prominently, spectra from the nitrogen-depleted condition (EMM-N) formed a distinctive cluster along the first LDA axis (Fig. 1D). Clusters of ethanol stress (EtOH 10%), carbon-source-depleted condition (EMM-C), glucose-limited condition (EMM 0.1% Glc.), heat-shock stress (Heat 39°C) and glucose-supplemented minimal medium (EMM 2% Glc.) were also well recognized by the LDA2-5 axes (Fig. 1E-G). Clusters from other conditions such as osmotic stress (Sorbitol 1 M), oxidative stress (H_2_0_2_ 2 mM), rich medium (YE) and heavy metal stress (CdS0_4_ 1 mM) mostly overlapped, so could not be recognized as separate clusters (Fig. 1G).

PC-LDA reports that the classification error is approximately 9.4%, i.e. test Raman spectra excluded from the calculation of the discriminatory bases are assigned to the correct cluster with 90.6% accuracy. Thus, clusters in the low-dimensional space reflect the characteristic differences of Raman spectra across conditions.

### Testing the linearity between Raman spectra and transcriptomes of *S. pombe*

We next asked whether the classification of Raman spectra in the low-dimensional space can be explained by other biological data. For this purpose, we obtained the transcriptomes of *S. pombe* cells under the same 10 culture conditions. All the transcripts including messenger RNAs (mRNA) and non-coding RNAs (ncRNA) except for ribosomal RNAs (rRNA) were annotated from PomBase [13, 14] (see Materials and Methods for details). Our hypothesis was that the transcriptome codes molecular compositions of the cell, and it linearly determines a low-dimensional Raman data **r**′*_ε_* obtained from cells in environment *ε*. In other words, process of estimating matrix **A** and predicting the Raman spectrum of the excluded condition for all environmental conditions.

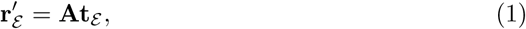

where **A** is a linear transformation matrix and **t**_*ε*_ is a 6560-dimension vector, in which each entry represents the expression level of a transcript in environment *ε*. **t**_*ε*_ are obtained from cell populations, not from single cells. Thus, we calculated the mean of single-cell Raman spectra from each environment *ε* to check the correspondence. To test the validity of this linear relation, we conducted a leave-one-out cross-validation (Fig. 2A). Out of all environmental conditions (10 conditions for *S. pombe),* we excluded one condition, and used the remaining 9 to estimate matrix **A** in Eq.(l). Matrix **A** was estimated by the partial least squares regression (PLS-R), which uniquely determines **A** from given datasets of **r**′*_ε_* and **t**′*_ε_* (see Materials and Methods) [15–17]. If our hypothesis is correct, one can predict the Raman spectrum of the excluded condition from the transcriptome data of its corresponding environment by Eq.(l) (Fig. 2A); the predicted Raman vector should be mapped onto the cluster of real spectra of the same environment. Changing the environmental condition to exclude, we repeated the same

**Figure 2.**
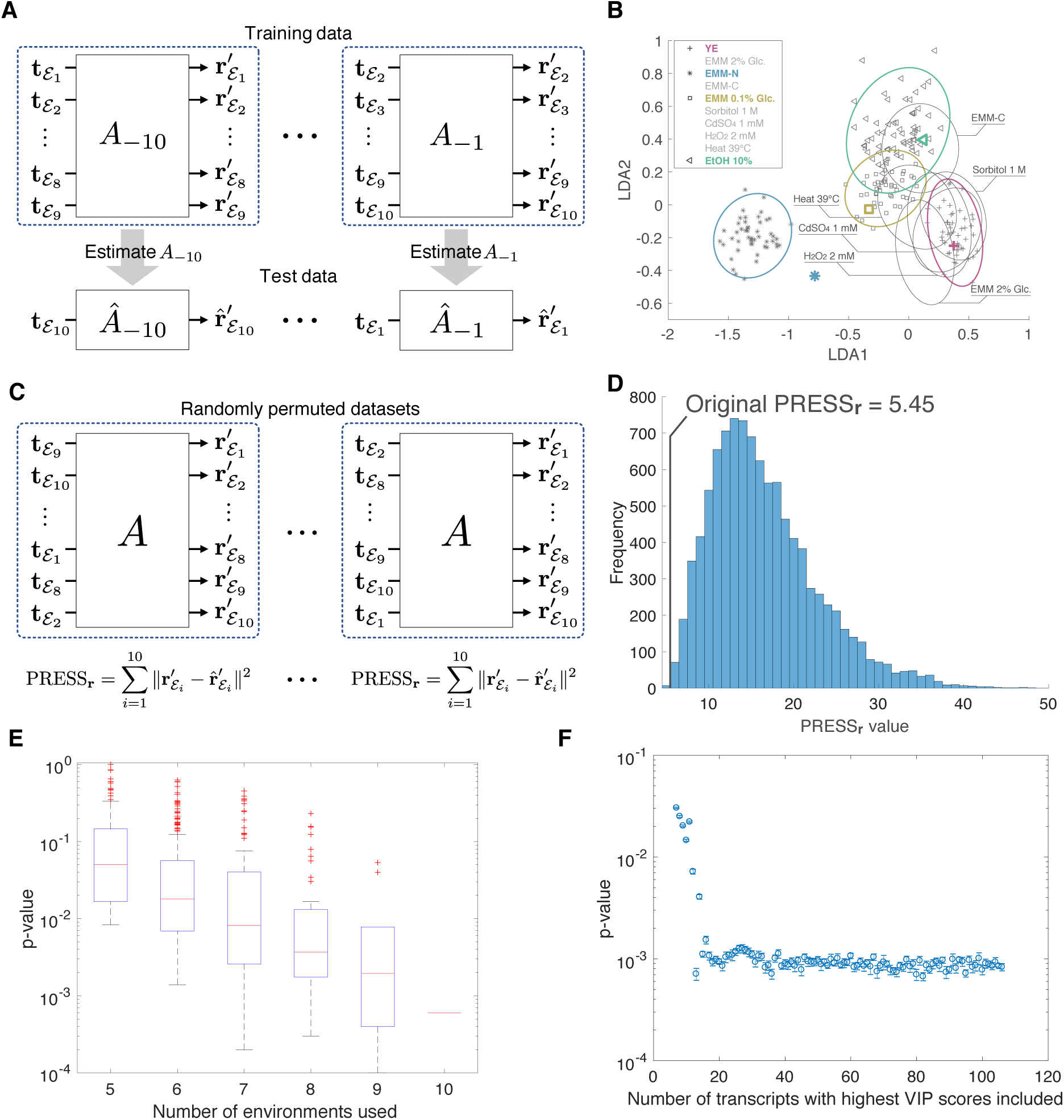
Linear correspondence between Raman spectra and transcriptomes. A. Leave-one-out crossvalidation. Out of all environmental conditions (*N* = 10 for *S. pombe),* one condition (ε_i_) is excluded, and the linear regression matrix **A**_–*i*_ is estimated by the partial least squares regression. Then, the excluded Raman spectrum is estimated by 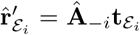. This is repeated for all *i* = 1, …,*N*. **B.** Example predictions of Raman spectra from the transcriptomes. Thick colored points represent Raman spectra predicted from the transcriptomes. C. Permutation test for significance of Raman-transcriptome linearity. 10,000 false datasets were created where environmental assignments of transcriptome data were randomly permuted. For each random permutation, PRESS_**r**_ was calculated and compared with the original PRESS_**r**_. **D.** Histogram of PRESS_**r**_ of 10,000 randomly permuted data. The original PRESS_r_ was 5.45, and the *p*-value was 0.0006. **E.** PRESS_**r**_ *p*-values when fewer numbers of environments were used. PRESS_**r**_ *p*-values were calculated for all possible combinations of environments for each number of environments (5-9 environments), and distributions of *p*-values were shown as box-and-whisker plots. **F.** *p*-values of PRESS_**r**_ when increasing the number of transcripts with highest VIP scores, *p*-values become stable after including 17 transcripts. The permutation test (10,000 permutations) was repeated 10 times per each point (error bar, standard error).

Our results show that predicted spectra were indeed assigned to positions within or adjacent to corresponding clusters (Fig. 2B and S3), supporting our hypothesis of linear correspondence.

To further test the reality of the observed linear correspondence, we calculated the predicted residual error sum of squares (PRESS) and compared it with randomized data. When estimating Â by PLS-R and using it to predict the Raman spectrum of the excluded condition, we calculated the prediction error defined as 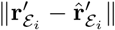, where **r**′_ε_i__ is the true data and 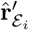 is the estimated data for environment *ε_i_*. We repeated the error calculation for all environments, and obtained the sum of squared errors

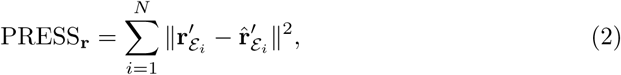

where *N* is the number of environmental conditions. In our case for *S. pombe, N* =10 and PRESS_r_ = 5.45.

When our hypothesis of linear correspondence is reasonable, PRESS_**r**_ should be small. To check this, we conducted the permutation test [18–20] by creating 10,000 false datasets in which environmental assignments of transcriptome data were randomly permuted (Fig. 2C). We calculated PRESS_**r**_ for these false data sets, and compared them to the original experimental value. We found that the original experimental PRESS_**r**_ was extremely small: the *p*-value of obtaining PRESS_**r**_ smaller than 5.45 was 0.0006 (Fig. 2D). This result offers strong support for the linear correspondence between Raman spectra and transcriptomes, and shows that the classification in the Raman space can be explained by differences of transcriptomes across conditions.

To further gain insight into the observed correspondence, we asked how many environments are necessary to find a good linear correspondence. To this end, we purposely selected fewer numbers of environments (5-9 environments), and calculated PRESS_r_ *p*-values for every combination of environments (Fig. 2E). For example, 252 possible combinations exist for 5 environments chosen among 10 (_10_C_5_) and 210 combinations for 6 environments chosen among 10 (_10_C_6_).

The result shows that *p*-values are generally smaller with more environments, and they all become smaller than 1% with 9 environments except for two combinations (Fig. 2E). These exceptional combinations lacked either EMM-N or EtOH 10%, a special environment distinguishable by axis LDA1 or LDA2 (Fig. 1D and 1E). These results show that having more environmental conditions and conditions in which cellular Raman spectra are largely distant from others, will generally improve the linear correspondence.

We next asked how many different kinds of transcripts are required to find a linear correspondence. To address this, we first evaluated the importance of each transcript based on the variable importance in projection (VIP) score in PLS-R analysis, which reflects the accumulated importance of each transcript to the linear regression [21, 22]. A high VIP score of a transcript indicates that its contribution to the linear correspondence is significant. The top 30 transcripts with the highest VIP scores are listed in Table 1. Then, starting from 7 transcripts (the minimum number of transcripts required to conduct PLS-R, see Materials and Methods for details) with the highest VIP scores, we increased the numbers of transcripts included and conducted the permutation test each time. Both *p*-values and PRESS_**r**_ values initially decreased, and plateaued after including 17 transcripts (Fig. 2F and S4). Thus, based on the VIP score, knowing the expression profiles of these 17 transcripts is sufficient to find a linear correspondence.

### Global expression profiles of *S. pombe* transcriptomes across conditions are predictable from Raman spectra

PLS-R not only estimates the linear transformation matrix **A**, but also conducts a dimension reduction of the transcriptome data. **W**e found that only 4 dimensions were required to explain 95% of the total variances of transcriptomes across conditions (Fig. 3A). The low-dimensionality of the transcriptome data indicates that global expression profiles of transcriptomes might also be predictable from Raman spectra. Note that this is a non-trivial inverse problem because we need to predict expression levels of 6560 transcripts in each environment only from 9-dimensional Raman data computed by PC-LDA.

**Figure 3.**
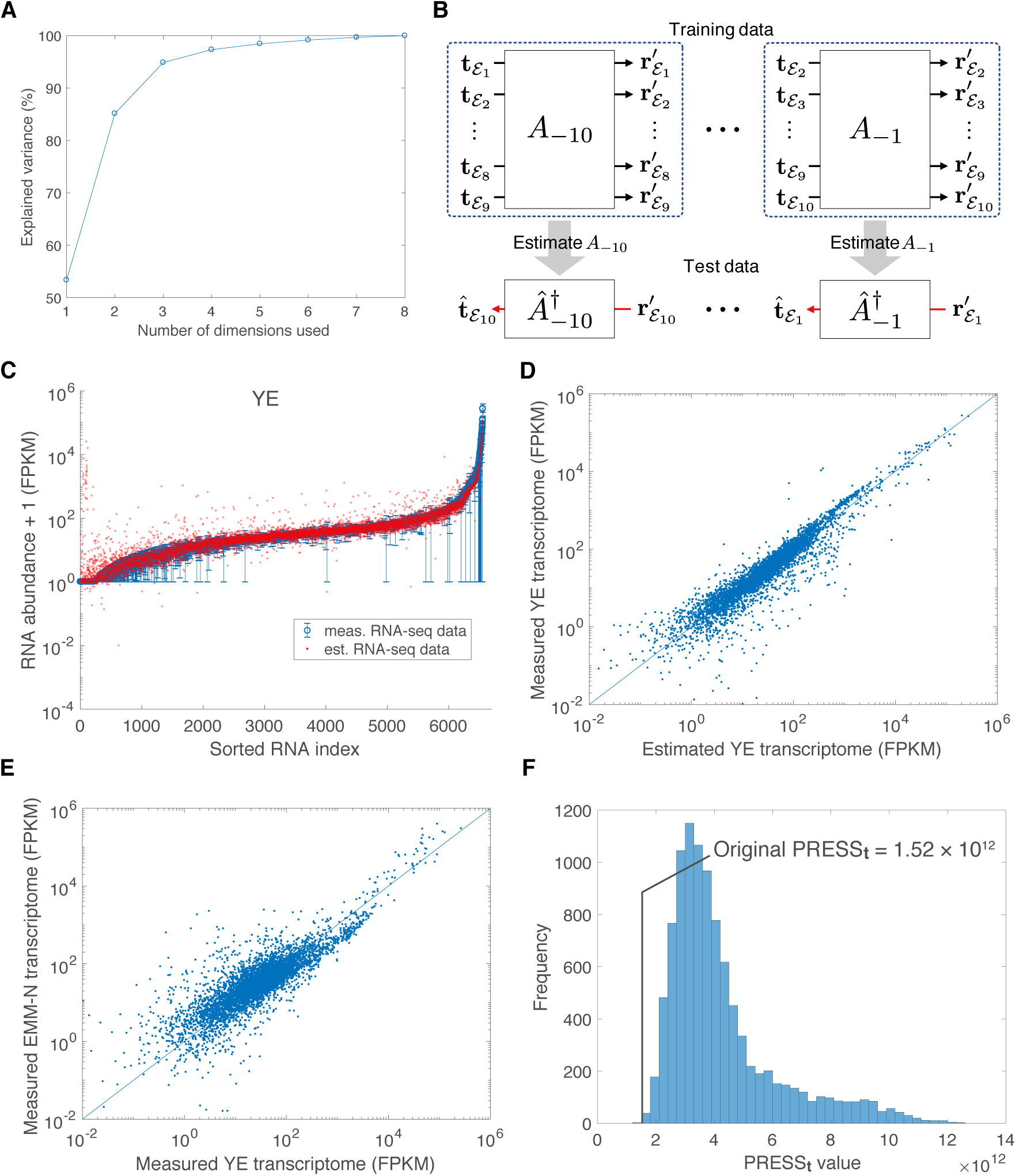
Predicting transcriptomes from Raman spectra. A. The total variance of transcriptome data explained by PLS-R. The horizontal axis represents the numbers of dimensions of transcriptomes after the dimension reduction by PLS-R used to calculate the variance, and the vertical axis the total variances explained. **B.** Predicting transcriptomes from Raman spectra. Transcriptomes were predicted by calculating the pseudo-inverse of Â_*–i*_ estimated in PLS-R as 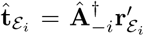. This is repeated for all *i* = 1,…,*N*. **C.** An example prediction of the transcriptome of rich medium environment (YE). Blue points represent the measured RNA abundance + 1 (FPKM) (average of the two replicate measurements; error bar, max-min range) sorted from low to high along the horizontal axis. Red points represent the RNA abundance predicted from Raman spectra. **D.** Scatter plot of the predicted YE medium transcriptome versus the measured YE medium transcriptome. **E.** Scatter plot of the measured YE medium transcriptome versus the measured nitrogen-depleted medium (EMM-N) transcriptome. **F.** Histogram of PRESS_**t**_ of 10,000 randomly permuted data. Environmental assignments of transcriptomes were randomly permuted 10,000 times, and PRESS_t_ were calculated for each permutation. The probability of accidentally finding PRESS_**t**_ less than the original experimental value, 1.52 × 10^12^, was extremely low (*p*-value 0.0004).

We estimated the global expression profile of transcriptome in environment *ε_i_* (denoted as 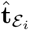) from obtained Raman spectra **r**′_*ε_i_*_. based on the linear relation of Eq.(l) and the Moore-Penrose pseudo-inverse of the PLS-R parameter Â_–*i*_ (Fig. 3B; see Materials and Methods for details). The results showed reasonably good agreements between 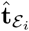 and **t**_ε_i__ (Fig. 3C, 3D, and S5). However, we also noticed that transcriptome data across conditions were already tightly correlated (Fig. 3E and S7), and likewise found relatively good agreements even between 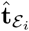 and transcriptome data of different environments (Fig. S8). We therefore evaluated in detail the precision level of our prediction by calculating the PRESS of transcriptomes,

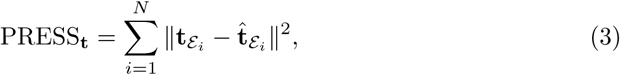

and again implemented the permutation test by randomly permuting the environmental correspondence between Raman and transcriptome data. Again for *S. pombe, N =* 10. Thereby we found that the original PRESS_**t**_ was very small: *p* = 0.0004 (Fig. 3F). The original prediction is thus significantly superior to randomly permuted data. In fact, transcriptomes predicted from Raman spectra explain the condition-dependent fold changes of mRNA and non-coding RNA transcripts (Fig. S6). These results prove that cellular Raman spectra allow us to capture the real global changes of the expression profile of transcriptomes across conditions in good precision. Note that without the low-dimensionality of transcriptomes, it is impossible to retrieve genome-wide expression profiles from the low-dimensional Raman spectra.

### The Raman-transcriptome correspondence in *E. coli*

To understand whether the observed Raman-transcriptome linearity is specific to *S. pombe* or more generally applicable to other organisms, we measured cellular Raman spectra and transcriptomes of *E. coli*. We focused on an *E. coli* strain MG1655 and its Δ*cyaA* mutant. *cyaA* encodes adenyl cyclase, which catalyzes the synthesis of cyclic AMP (cAMP) from ATP [23]. The growth of the Δ*cyaA* mutant is suppressed in cAMP-depleted culture media, but restored by exogenous cAMP supplement in a concentration-dependent manner [24]. We measured cellular Raman spectra (Fig. 4A, 4B and S9) and transcriptomes of Δ*cyaA* mutant cultured in the media with 0, 0.1, 0.5 and 1 mM cAMP, and those of the parental MG1655 strain (no cAMP in the medium) all sampled from late-exponential phase (OD_660_ = 0.8).

**Figure 4.**
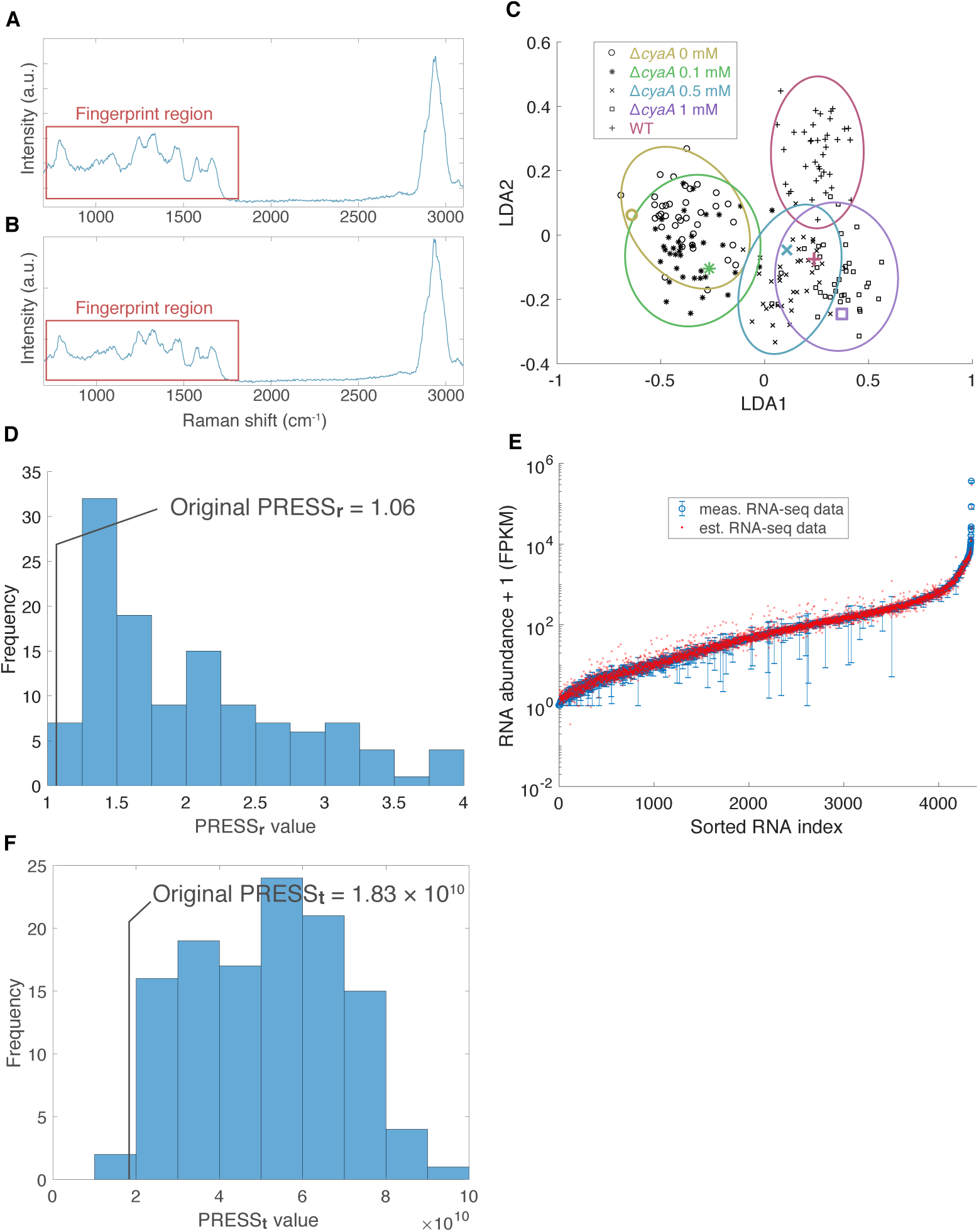
Raman-transcriptome correspondence in *E. coli*. A, B. Example Raman spectra of *E. coli* for wild type (**W**T, plot A), and Δ*cyaA* mutant supplemented with 0 mM cAMP Δ*cyaA* 0 mM, plot B). **R**ed rectangles represent the fingerprint region. **C.** Dimension reduction of *E. coli* Raman spectra by PC-LDA. Black points represent the measured Raman spectra after dimension reduction by PC-LDA shown on the LDA1-LDA2 plane. Colored ellipses represent the χ^2^ 95% confidence intervals for different culture conditions: Yellow for Δ*cyaA* 0 mM; green for Δ*cyaA* 0.1 mM; blue for Δ*cyaA* 0.5 mM; purple for Δ*cyaA* 1 mM; red for WT. Thick colored points denote the low-dimensional Raman data predicted from the corresponding transcriptomes. **D.** PRESS_r_ histogram when randomly permuting environmental assignments of transcriptome data. The original experimental PRESS_r_ = 1.06 was the third lowest of all 5! = 120 possible permutations (*p*-value = 3/120 = 0.0250). E. Example prediction of *E. coli ΔcyaA* 0 mM transcriptomes from Raman spectra by 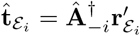. Blue points represent the measured RNA abundance + 1 (FPKM) (average of the three replicate measurements; error bar, standard error) sorted from low to high along the horizontal axis. Red points represent the RNA abundance predicted from Raman spectra. **F**. PRESS_**t**_ histogram when randomly permuting environmental assignments of transcriptome data. The original experimental PRESS_**t**_ = 1.83 × 10^10^ was the second lowest of all 5! = 120 possible permutations (*p*-value = 2/120 = 0.0167).

As was done for *S. pombe,* we first conducted PC-LDA to obtained Raman spectra, finding that spectra under five different conditions (4 for the Δ*cyaA* mutant, and 1 for wild type) can be classified in a low-dimensional Raman space (Fig. 4C). Interestingly, clusters of Raman spectra of the mutant became closer to that of wild type as the concentration of exogenous cAMP increased. Next we conducted PLS-R to find a linear regression, and found that Raman spectra predicted from transcriptome data were assigned within or adjacent to the correct clusters (Fig. 4C). We calculated PRESS_**r**_ by Eq. (2) where *N* = 5 for *E. coli,* and subsequently conducted a permutation test. The test showed that the original combination of the Raman and transcriptome data gave the third lowest PRESS_**r**_ (1.06) among the 5!(= 120) possible permuted combinations (PRESS_**r**_ *p*-value 0.0250, Fig. 4D). The prediction of transcriptomes from Raman spectra also gave good results, giving the second smallest PRESS_**t**_ (PRESS_**t**_ *p*-value 0. 0167, Fig. 4E, 4F, Sil, S10-S13). Together, this confirms the linear correspondence between Raman spectra and transcriptomes even in *E. coli,* and indicates that it should have broader applicability to other organisms and cell types.

### ncRNAs are largely responsible for the Raman-transcriptome correspondence in *S. pombe*

We next examined what types of transcripts were responsible for establishing the observed Raman-transcriptome linear correspondence. Based on the list of transcripts sorted by the VIP scores, we found out that the main contributors were largely ncRNAs including small nucleolar RNAs (snoRNAs) and tRNAs in *S. pombe* (Table 1): The highest scoring mRNA was ranked only at the 55-th from the top.

**Table 1.** List of *S. pombe* transcripts with top 30 VIP scores. The VIP score of each transcript is shown on the third column. The mean expression level of each transcript across 10 environmental conditions is shown on the fourth column, in the unit of fragments per kilobase per million mapped reads (FPKM). Type “N” on the fifth column indicates that the transcripts are known or predicted non-coding RNAs.

| ID | Name | VIP | FPKM | Type | Description |
| --- | --- | --- | --- | --- | --- |
| SPSNRNA.06 | snu6 | 52.3 | $3.95 \times 10^5$ | N | small nuclear RNA U6 |
| SPNCRNA.98 | srp7 | 26.6 | $3.86 \times 10^5$ | N | 7SL signal recognition particle component |
| SPMITTRNALYS.01 | SPMITTRNALYS.01 | 23.1 | $1.81 \times 10^5$ | N | tRNA Lysine, mitochondrial |
| SPNCRNA.510 | SPNCRNA.510 | 21.1 | $1.46 \times 10^5$ | N | non-coding RNA (predicted) |
| SPSNORNA.24 | snoR39b | 18.8 | $1.32 \times 10^5$ | N | small nucleolar RNA R39b (predicted) |
| SPSNORNA.31 | snoR39a | 12.1 | $7.27 \times 10^4$ | N | small nucleolar RNA snR39 |
| SPSNORNA.43 | snR91 | 11.2 | $1.29 \times 10^5$ | N | box H/ACA small nucleolar RNA snR91 |
| SPSNORNA.17 | snoR58 | 11.2 | $7.54 \times 10^4$ | N | small nucleolar RNA snR58 (predicted) |
| SPMITTRNAGLY.01 | SPMITTRNAGLY.01 | 10.5 | $6.44 \times 10^4$ | N | tRNA Glycine, mitochondrial |
| SPSNORNA.32 | sno12 | 10.4 | $1.29 \times 10^5$ | N | box H/ACA small nucleolar RNA 12/snR99 |
| SPSNORNA.13 | snoR69b | 9.87 | $9.17 \times 10^4$ | N | small nucleolar RNA snoR69b (predicted) |
| SPSNORNA.27 | snoR47 | 9.25 | $5.38 \times 10^4$ | N | small nucleolar RNA R47 (predicted) |
| SPMITTRNAALA.01 | SPMITTRNAALA.01 | 8.25 | $2.88 \times 10^4$ | N | tRNA Alanine, mitochondrial |
| SPSNORNA.16 | snoR56 | 8.03 | $6.96 \times 10^4$ | N | small nucleolar RNA snR56 (predicted) |
| SPNCRNA.507 | SPNCRNA.507 | 7.88 | $7.79 \times 10^4$ | N | non-coding RNA (predicted) |
| SPSNORNA.01 | snR40 | 7.80 | $4.02 \times 10^4$ | N | small nucleolar RNA snR40 (predicted) |
| SPATRNAILE.02 | SPATRNAILE.02 | 7.77 | $3.10 \times 10^4$ | N | tRNA Isoleucine |
| SPMITTRNATYR.01 | SPMITTRNATYR.01 | 7.42 | $2.46 \times 10^4$ | N | tRNA Tyrosine, mitochondrial |
| SPMITTRNAASP.01 | SPMITTRNAASP.01 | 7.41 | $3.10 \times 10^4$ | N | tRNA Aspartic acid, mitochondrial |
| SPSNORNA.21 | snoU14 | 6.97 | $5.77 \times 10^4$ | N | small nucleolar RNA U14 |
| SPSNORNA.41 | snR46 | 6.29 | $5.33 \times 10^4$ | N | box H/ACA small nucleolar RNA snR46 |
| SPBTRNAARG.04 | SPBTRNAARG.04 | 5.98 | $2.61 \times 10^4$ | N | tRNA Arginine |
| SPMITTRNAGLU.01 | SPMITTRNAGLU.01 | 5.52 | $2.39 \times 10^4$ | N | tRNA Glutamic acid, mitochondrial |
| SPCTRNAARG.08 | SPCTRNAARG.08 | 5.46 | $2.12 \times 10^4$ | N | tRNA Arginine |
| SPBTRNAGLY.09 | SPBTRNAGLY.09 | 5.43 | $1.53 \times 10^4$ | N | tRNA Glycine |
| SPSNRNA.01 | snu1 | 5.27 | $6.80 \times 10^4$ | N | small nuclear RNA U1 |
| SPSNORNA.44 | snR92 | 4.80 | $5.22 \times 10^4$ | N | box H/ACA small nucleolar RNA snR92 |
| SPSNRNA.07 | snu32 | 4.76 | $5.78 \times 10^4$ | N | small nucleolar RNA U3B |
| SPBTRNATHR.06 | SPBTRNATHR.06 | 4.71 | $1.89 \times 10^4$ | N | tRNA Threonine |
| SPATRNAVAL.01 | SPATRNAVAL.01 | 4.56 | $1.20 \times 10^4$ | N | tRNA Valine |

To further understand this result, we separated transcriptomes of *S. pombe* into mRNAs (containing 5091 transcripts) and ncRNAs (containing 1469 transcripts), and checked whether a linear correspondence can be found between Raman spectra and these coding and non-coding subsets of transcriptomes (Fig. 5A). The result showed that the linear correspondence between Raman spectra and mRNAs was as poor as randomly permuted data (PRESS_r_mRNA__ *p*-value = 0.4803), whereas the correspondence with the ncRNA subset was excellent (PRESS_r_ncRNA__ *p*-value = 0.0009, Fig. 5A). This also confirms the importance of ncRNAs to find the linear correspondence between Raman spectra and transcriptomes.

**Figure 5.**
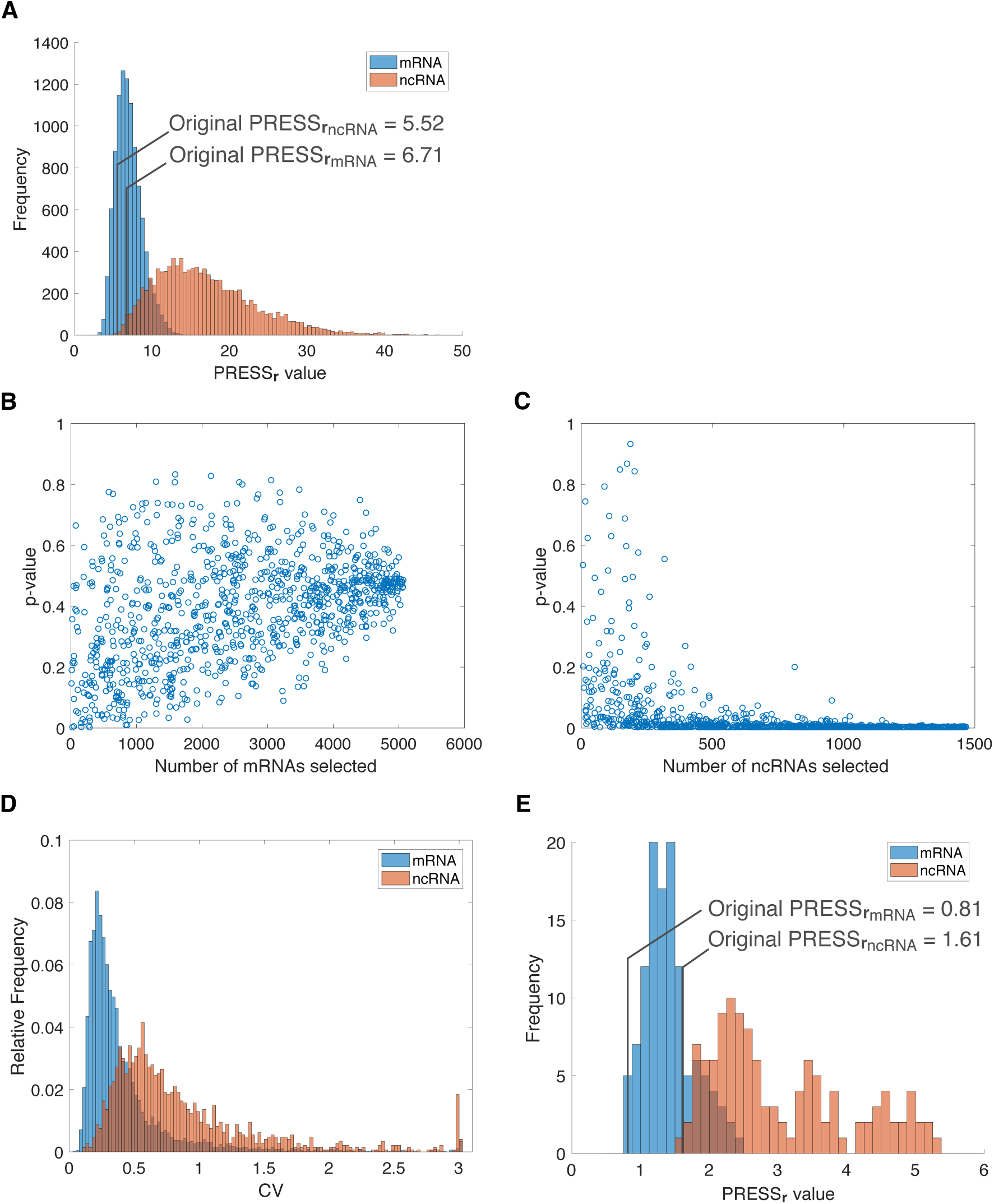
Transcriptome subsets contributing to the Raman-transcriptome correspondence. A. PRESS_**r**_ ahistograms of randomly permuted data sets of *S. pombe* mRNAs (blue) and ncRNAs (red). Original PRESS_**r**_ of mRNA was 6.71 (*p*-value, 0.4803) and that of ncRNAs was 5.52 (*p*-value, 0.0009). **B.** PRESS_**r**_ *p*-values of randomly selected mRNA subsets in *S. pombe*. 1,000 subsets were selected, and 1,000 random permutations per each subset were conducted to calculate *p*-values. C. PRESS_**r**_ *p*-values of randomly selected ncRNA subsets in *S. pombe*. 1,000 subsets were selected, and 1,000 random permutations per each subset were conducted to calculate *p*-values. In stark contrast to mRNAs, the vast majority of ncRNA subsets had small *p*-values. **D.** Histograms of coefficient of variations of mRNAs (blue) and ncRNAs (red) across 10 environmental conditions in *S. pombe*. **E.** PRESS_**r**_ histograms of randomly permuted data sets of *E. eoli* mRNAs (blue) and ncRNAs (red). Original PRESS_**r**_ of mRNA was 0.81 (*p*-value, 0.0250) and that of ncRNAs was 1.61 (*p*-value, 0.0167).

We also randomly sampled different numbers of transcripts either from the mRNA or ncRNA subset, and searched for the presence of a linear correspondence between those randomly sampled subsets and Raman spectra. Our results show that only very limited combinations of the sampled mRNA subsets yielded linearity (Fig. 5B), but many subsets of ncRNAs could correspond even with small numbers of transcripts (Fig. 5C). This result again indicates the superiority of ncRNAs for finding a linear correspondence.

Note that these results do not indicate that Raman spectra cannot predict mRNA profiles: our prediction of mRNA expression levels from Raman spectra was actually more precise than ncRNA expression levels as below. To evaluate the prediction accuracy of transcriptomes, we calculated the “coefficient of variation of prediction error per each transcript” as follows:

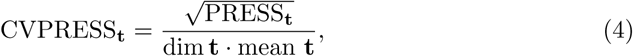

where dim **t** is the total number of transcripts, and mean **t** is the mean expression level of all transcripts in **t** across all environment conditions. Here, we used **t**_mRNA_ or **t**_ncRNA_ for **t**. **W**e found that CVPRESS_t_mRNA__ = 0.0909 and CVPRESS_t_ncRNA__ = 0.383, showing that the prediction accuracy of each mRNAs relative to their mean expression levels was actually higher than that of ncRNAs. This may be counterintuitive to the fact that PRESS_r_mRNA__ *p*-value is high. Instead, this indicates that expression levels of mRNAs do not change much across conditions, and there is not much difference even when the data set is randomly permuted. In fact, the coefficient of variations (CV) of expression levels across conditions were larger for ncRNAs than for mRNAs (Fig. 5D). Also, PRESS_r_ncRNA__ of randomized data changed over a much broader range than mRNAs (Fig. 5A), showing once again that expression levels of ncRNAs change more dynamically in response to environmental changes.

We likewise conducted the same analysis for *E. coli*. In *E. coli,* the number of ncRNAs is much smaller than that of *S. pombe,* constituting only 2.18% of the *E. coli* transcriptome. The analysis revealed that the linear correspondence can be found with both mRNA and ncRNA subsets (*p*-value = 0.0250 for mRNA, and 0.0167 for ncRNA, Fig. 5E). In fact, 28 among the 30 top VIP-scored transcripts were mRNAs (Table 2). Random sampling test also showed that the linear correspondence was more easily found with mRNA subsets than with ncRNA subsets for the current 5 conditions (Fig. S14). Therefore, the necessity of ncRNAs for the Raman-transcriptome linearity was not as apparent as that in *S. pombe*.

**Table 2.** List of *E. coli* transcripts with top 30 VIP scores. The VIP score of each transcript is shown on the third column. The mean expression level of each transcript across 5 environmental conditions is shown on the fourth column, in the unit of FPKM. Type “N” and “C” on the fifth column indicate ncRNAs and mRNAs, respectively.

| ID | Name | VIP | FPKM | Type | Description |
| --- | --- | --- | --- | --- | --- |
| b2621 | ssrA | 54.3 | $2.57 \times 10^5$ | N | tmRNA, 10Sa RNA |
| b3123 | rnpB | 21.9 | $8.98 \times 10^4$ | N | RNase P, M1 RNA component |
| b1677 | lpp | 14.2 | $2.94 \times 10^4$ | C | murein lipoprotein |
| b2579 | grcA | 13.5 | $9.88 \times 10^3$ | C | autonomous glycyl radical cofactor |
| b3510 | hdeA | 8.98 | $1.73 \times 10^4$ | C | stress response protein acid-resistance protein |
| b3708 | tnaA | 5.98 | $2.90 \times 10^3$ | C | tryptophanase/L-cysteine desulfhydrase, PLP-dependent |
| b2266 | elaB | 5.64 | $7.85 \times 10^3$ | C | putative membrane-anchored DUF883 family ribosome-binding protein |
| b3556 | cspA | 4.28 | $1.04 \times 10^4$ | C | RNA chaperone and antiterminator, cold-inducible |
| b3985 | rplJ | 3.32 | $9.30 \times 10^3$ | C | 50S ribosomal subunit protein L10 |
| b4217 | ytfK | 3.05 | $4.27 \times 10^3$ | C | DUF1107 family protein |
| b2215 | ompC | 2.93 | $1.48 \times 10^4$ | C | outer membrane porin protein C |
| b3986 | rplL | 2.90 | $8.53 \times 10^3$ | C | 50S ribosomal subunit protein L7/L12 |
| b0953 | rmf | 2.88 | $3.24 \times 10^3$ | C | ribosome modulation factor |
| b0812 | dps | 2.85 | $4.45 \times 10^3$ | C | Fe-binding and storage protein; stress-inducible DNA-binding protein |
| b3314 | rpsC | 2.66 | $5.94 \times 10^3$ | C | 30S ribosomal subunit protein S3 |
| b2096 | gatY | 2.65 | $2.01 \times 10^3$ | C | D-tagatose 1,6-bisphosphate aldolase 2, catalytic subunit |
| b0957 | ompA | 2.65 | $1.35 \times 10^4$ | C | outer membrane protein A (3a;II*;G;d) |
| b3316 | rpsS | 2.64 | $5.82 \times 10^3$ | C | 30S ribosomal subunit protein S19 |
| b2343 | yfcZ | 2.60 | $3.88 \times 10^3$ | C | UPF0381 family protein |
| b3296 | rpsD | 2.57 | $7.45 \times 10^3$ | C | 30S ribosomal subunit protein S4 |
| b3315 | rplV | 2.51 | $5.68 \times 10^3$ | C | 50S ribosomal subunit protein L22 |
| b3509 | hdeB | 2.50 | $5.08 \times 10^3$ | C | acid-resistance protein |
| b4240 | treB | 2.48 | $1.32 \times 10^3$ | C | trehalose-specific PTS enzyme: IIB and IIC component |
| b3307 | rpsN | 2.47 | $6.91 \times 10^3$ | C | 30S ribosomal subunit protein S14 |
| b2092 | gatC | 2.46 | $1.65 \times 10^3$ | C | pseudogene, galactitol-specific enzyme IIC component of PTS; transport; Transport of small molecules: Carbohydrates, organic acids, alcohols; PTS system galactitol-specific enzyme IIC |
| b0814 | ompX | 2.44 | $5.12 \times 10^3$ | C | outer membrane protein X |
| b3308 | rplE | 2.42 | $7.31 \times 10^3$ | C | 50S ribosomal subunit protein L5 |
| b3065 | rpsU | 2.40 | $7.17 \times 10^3$ | C | 30S ribosomal subunit protein S21 |
| b3319 | rplD | 2.39 | $6.02 \times 10^3$ | C | 50S ribosomal subunit protein L4 |
| b4015 | aceA | 2.38 | $3.12 \times 10^3$ | C | isocitrate lyase |

## Discussion

Cellular Raman spectra reflect the comprehensive molecular compositions of cells, and therefore spectral differences should be associated with cellular state differences. However, interpreting spectra has been challenging due to the difficulty of decomposing total cellular spectra into those of constituent biomolecules. In this report, instead of pursuing the spectral decomposition, we asked whether whole-cell Raman spectra could be directly and computationally corresponded to other types of well studied omics-level information. Employing dimension reduction methods, we showed that dimensions of high dimensional cellular Raman spectra and transcriptomes measured by RNA-seq can be greatly reduced, and connected linearly through a shared low-dimensional subspace. Accordingly, we were able to reconstruct global gene expression profiles by applying the calculated transformation matrix to cellular Raman spectra, and vice versa. We therefore provided firm experimental evidence that the differences of cellular Raman spectra contain key information that allows us to detect cellular states.

The linear correspondence between cellular Raman spectra and transcriptomes is far from trivial because transcriptomes targeted in our study (the total RNA excluding rRNAs) constitute only a small fraction of biomass: the total RNA excluding rRNAs constitute 2% of the biomass, whereas proteins constitute 40-50% in *Saccharomyces cerevisiae* [25, 26]. Furthermore, Raman signals mostly come from proteins, and the contribution of total RNAs to the total signal is considered minor [27, 28]. It is thus implausible that the observed Raman spectra directly reflects signals from RNAs targeted in our study. The observed linear correspondence instead indicates that whole-cell molecular compositions of cells are tightly and linearly constrained by the transcriptome. Cellular Raman signals come from all the constituent biomolecules in a cell including proteins, lipids, and metabolites, but our PLS-R analysis did not take into account of the abundances of biomolecules other than a fraction of RNAs. The fact that Raman spectra and transcriptomes correspond linearly implies that abundances of other biomolecules might also be linearly related to the transcriptome. This speculation could be tested by trans-omics analyses to examine the correspondences among multi-level omics data such as proteomes, metabolomes, and transcriptomes [29]. The unexpected correspondence between Raman spectra and transcriptomes might also imply that similar multivariate analyses could find linkages between other types of non-destructive spectroscopic data such as whole-cell NMR [30, 31] and omics data. It would be an important subject to explore such technical possibilities for future cell analysis study.

It is also intriguing that in *S. pombe,* ncRNAs are more linearly corresponded to Raman spectra than are mRNAs because ncRNAs do not contribute to proteomes directly. Our analysis reveals that snoRNAs and tRNAs had high VIP scores in *S. pombe* (Table 1). These ncRNAs are directly or indirectly involved in translational processes: For example, snoRNAs are known to be necessary for the maturation of ribosomal RNAs [32, 33]. Therefore, expression levels and the combined action of these ncRNAs may influence the translation of all proteins and consequently modulate global chemical composition of cells. Importantly, our results indicate that changes of mRNA expression levels in *S. pombe* across environment conditions are subtle relative to ncRNAs (Fig. 5D), which might have prevented us from finding a linear correspondence between Raman spectra and mRNA transcriptome subset. Note that our results do not exclude the possibility that mRNA profiles are linked to Raman spectra non-linearly.

On the other hand, ncRNAs are much less abundant in *E. coli,* and most of the transcripts with high VIP scores were mRNAs (Table 2). However, we found that the VIP list contained many transcripts coding ribosomal subunit proteins and ribosome modulating factors (Table 2). Taken together, our findings might indicate that alterations in translation machinery is one of the major cellular responses to environmental changes, and thus intimately linked to cellular global molecular compositions that are reflected in Raman spectra.

Our results showing that the transcriptome is low-dimensional indicate that intracellular gene expression is globally coupled and that expression-level changes of many genes occur in a coordinated manner. The degree of freedom in transcriptomes should therefore be severely limited, which has been in fact suggested in many microarray and sequencing studies [34–43]. Importantly, as shown in our study, such global changes of transcriptomes are associated with changes of cellular Raman spectra, which can now be monitored non-destructively at the single-cell level and in a snapshot manner. Furthermore, if the transformation parameter is known beforehand, one could estimate instantly the change of expression levels of each transcript from Raman spectra as conducted in Fig. 3. It should be noted that the low-dimensionality of transcriptomes was indispensable for our prediction because they were predicted based on dimension-reduced Raman spectra; changes of transcriptomes that require more dimensions than the total number of LDA axes are unpredictable in principle. Therefore, the fact that we could predict transcriptomes in a reasonably good precision in turn provides evidence for the low-dimensionality of transcriptomes.

Single-cell Raman microscopy is compatible with live-cell time-lapse imaging, though the photo-damage on cells by incident laser and background spectral noise from culture media must be carefully considered. Our results therefore indicate that single-cell Raman spectra have the potential to provide omics information directly from living cells in a non-destructive and snapshot manner. Such *spectroscopic live-cell omics* studies would provide the way to investigate how global cellular states dynamically change in single living cells across diverse environmental conditions and cell types. If the Raman-transcriptome correspondence is further confirmed for other cell types, single-cell Raman microscopy could be applied to detecting distinct cells such as malignant cancers, pluripotent stem cells and antibiotic-resistant bacteria.

## Materials and Methods

### *S. pombe* strain and culture conditions

A haploid strain, 972 h^−^, was used for all *S. pombe* experiments. Initially, cells were cultured at 30°C in Yeast Extract (YE) (Bacto Yeast Extract (Becton Dickinson and Co)) + 3% glucose liquid medium until OD_600_ =0.7-1.0. Then, cell cultures were inoculated to the following stress conditions in liquid cultures: YE + 1 mM CdS0_4_, YE + 1 M Sorbitol, YE + 2 mM H_2_0_2_, heat shock in a water bath at 39°C for an hour. For the other stress conditions, cell cultures were washed three times with the media EMM-N, EMM-C, EMM 2% Glucose, EMM 0.1% Glucose, YE + 10% EtOH (Table S1), and cultured in each medium at 30°C for 24 hours.

Prior to Raman microscopy measurements, cells from each stress condition were washed with phosphate buffered saline (PBS) three times, and fixed with 2% formaldehyde in PBS for an hour at 4°C. Then, cells were washed once with PBS + 100 mM glycine to quench the free aldehyde, and twice with PBS. Subsequently, all samples were stored at 4°C until they were measured.

### *E. coli* strains and culture conditions

*E. coli* MG1655 Δ*cyaA* strain was constructed by P1 transduction from BW25113 Δ*cyaA* strain in Keio collection [44]. The deletion of *cyaA* ORF was verified after isolation by genome sequencing.

*E. coli* MG1655 strain was cultured in 10 mL L-broth (1.0% Bacto Tryptone (Becton Dickinson and Co.), 0.5% Bacto Yeast Extract (Becton Dickinson and Co.) and 0.5% NaCl). MG1655 Δ*cyaA* strain was cultured in 10 mL L-broth containing 0, 0.1, 0.5 or 1.0 mM cAMP. Cells were grown at 37°C to late exponential phase (OD_660_=0.8). Prior to Raman microscopy measurements, cells were washed twice with physiological saline.

### Raman microscopy

Raman spectra of cells were obtained with a custom-built Raman microscope where a commercial Raman imaging system (STR-Raman, AIRIX corp.) was integrated into a Nikon Ti-E microscope. A 532 nm, continuous-wave diode-pumped solid-state laser (Gem 532, Laser Quantum) was used as excitation. For *S. pombe,* a 60x/NA 1.2 water immersion objective lens (Olympus, UPLSAPO 60XW) was used at 4 mW power at the sample stage. For *E. coli,* a 100×/NA 0.9 air objective lens (Olympus, MPLN 100X) was used at 18 mW power. Backscattered Raman signals were focused through a 100 *μ*m pinhole, dispersed by a spectrometer (Acton SP2300i, Princeton Instruments) equipped with a 300 gr/mm grating, and detected with a sCMOS camera (Orca Flash 4.0 v2, Hamamatsu Photonics). To reduce dark noise, the sCMOS camera was water-cooled at 15°C. The exposure time of each cell was 10 seconds.

Unlike CCD detectors, sCMOS detectors have pixel-dependent readout noise that must be reduced for actual use in Raman microscopy. To address this, a sCMOS specific noise reduction filter inspired by [45] was implemented. All of the following analyses were conducted by scripts written in Matlab 2017a. First, 10, 000 blank, 2048 × 2048 pixel images with exposure time of 10 seconds were obtained to characterize the noise distribution of each pixel. The offset *o_i_* and variance var_i_ of pixel *i* were calculated as follows:

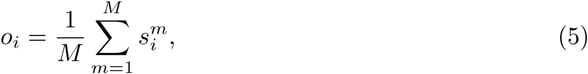

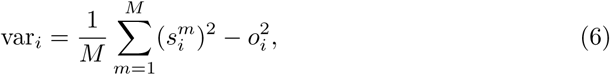

where *M* is the total number of images taken (in this case *M* = 10,000), *m* is the frame number of obtained dark images and 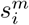 the analog-to-digital unit (ADU) count of pixel *i* at frame *m*. An offset subtraction for each pixel was conducted, and a 2-dimensional convolution filter was applied to obtained images as follows:

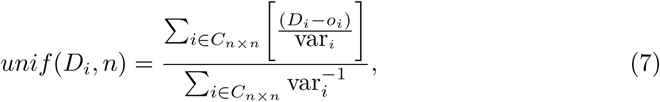

where *D_i_* is the ADU count of each pixel, *n* is the number of pixels per window size and *C_n×n_* represents the kernel region which is a *n* × *n* square box centered around pixel *i*. In our study, *n* = 3. In essence, this convolution filter assigns a weighted average to pixel counts, where pixels with low variances are given higher weights than those with high variances. After applying the convolution filter, the region of the spectrum in the image was cropped, and the sum of pixel counts along the direction perpendicular to the wavenumber was calculated to obtain a Raman spectrum. The wavenumber was calibrated referencing the standard Raman spectrum of ethanol, and spectral regions of 632 to 1873 cm^−1^ was used for all subsequent multivariate analyses (2 cm^−1^ per pixel). Furthermore, each spectrum was smoothed by the Savitzky-Golay filter [46], and normalized by subtracting the mean and dividing it by its standard deviation.

For preparing *S. pombe* samples for Raman measurements, 1 *μL* of cell suspension was placed on a synthetic quartz slide glass put in place with a synthetic quartz cover glass (TOSHIN RIKO CO., LTD). The rims of the coverslips were sealed with Vaseline to prevent evaporation during measurements. The center of 15-26 cells were measured for every sample, and three biological replicates were obtained, which resulted in a total of 54-76 cell measurements per each environment condition.

For *E. coli,* 5 *μ*L of cell suspension was placed on a synthetic quartz slide glass, and air dried for 5-10 minutes. 15 cells were measured for every sample, and three biological replicates were obtained, which resulted in a total of 45 cell measurements per each condition. 5 background spectra for each slide glass were obtained, and the average was subtracted from obtained cellular spectra.

### *S. pombe* RNA sequencing and data processing

For *S. pombe* RNA-seq, two biological replicates were measured. 50 mL cell cultures of each environmental condition were prepared as described above. Each culture was pelleted down, and quickly frozen with liquid nitrogen. The total RNA was extracted as described in [47], followed by a DNA removal (RQ1 DNase, Promega) and a ribosome RNA removal by the Ribo-Zero Gold rRNA Removal kit for yeast (Illumina Inc.). Sequencing libraries were prepared using NEBNext Ultra Directional RNA Library Prep Kit for Illumina (NEB) following the manufacturer’s instructions. 150 bp paired-end sequencing was conducted with MiSeq (Illumina Inc.). Raw RNA-seq reads were mapped on to the reference genome of 972 h^−^ *S. pombe* (ASM294v2) from PomBase [13, 14] by TopHat2 [48], and FPKMs of annotated genes and noncoding transcripts were calculated by Cufflinks 2.0 [49, 50].

### *E. coli* RNA sequencing and data processing

For *E. coli* RNA-seq, three biological replicates were measured. In the preparation of RNAs, RNAprotect Bacteria Reagent (Qiagen N.V.) was added to exponential phase cultures, and then, cells were lysed using lysozyme (SEIKAGAKU Co.). Then, the total RNA was extracted from the lysates using an RNeasy mini kit (Qiagen N.V.) and RNase-free DNase set (Qiagen N.V.) following the manufacturer’s instructions. Sequencing libraries were prepared by the NEBNext mRNA library prep kit for Illumina (NEB) with the following modifications. The random hexamer primer was used for reverse transcription. After second strand synthesis, double stranded cDNA was fragmented to an average length of 300 bp using a Covaris S2 sonication system (Covaris Inc.). One hundred cycles of paired-end sequencing were carried out using HiSeq 2500 system (Illumina Inc.) following the manufacturer’s instructions. After the sequencing reactions were complete, the Illumina analysis pipeline (CASAVA 1.8.0) was used to process the raw sequencing data. RNA-seq reads were trimmed using CLC Genomics Workbench ver. 8.5.1 (Qiagen N.V.) with the following parameters; Phred quality score >30; Removing terminal 15 nucleotides from 5’ end and 3 nucleotides from 3’ end; Removing truncated reads less than 30 nucleotides length. Trimmed reads were mapped to all genes in *E. coli* strain MG1655 (accession number: NC_000913.3) using CLC Genomics Workbench ver. 8.5.1 (Qiagen N.V.) with the following parameters; Length fraction: 0.7; Similarity fraction: 0.9; Maximum number of hits for a read: 1. The expression level of each gene was calculated by counting the mapped reads to each gene and were normalized by calculating the values of FPKM. All transcripts were annotated from [51].

### Principal component-linear discriminant analysis (PC-LDA)

To reduce systematic-error and dimensions of cellular Raman spectra, we conducted principal component-linear discriminant analysis (PC-LDA). In short, PC-LDA is a supervised classification technique that combines principal component analysis (PCA) and linear discriminant analysis (LDA) to find the most discriminatory bases while avoiding over-fitting. PCA is first applied to the original high-dimensional Raman spectra to reduce noise and dimension, which simultaneously reduces over-fitting and enables conducting the following LDA analysis against high-dimensional data [7, 8, 12]. In our study, for both *S. pombe* and *E. coli,* we used principal components that in total explained 98% of the variance of the original Raman spectra. Then, against the chosen principal components, LDA takes into account the culture-condition assignments and extracts the most discriminatory bases by maximizing the ratio of the between-group variance to the sum of within-group variances in the lower dimensional space.

To test how well PC-LDA was able to classify cellular Raman spectra, 1/6-th of the spectra from every environment condition was excluded from the calculation of the discriminatory bases, projected on to the PC-LDA space, and classified by the maximum likelihood method. It was assumed that single-cell Raman spectra in the PC-LDA space from each environment followed a Gaussian distribution, and the excluded Raman spectra were classified as the environment which gave the highest likelihood. For *S. pombe,* the classification accuracy was 90.6%, and for *E. coli,* 65.7%.

### Prediction of Raman spectra and transcriptomes by partial least squares regression (PLS-R)

To evaluate the linearity between dimension reduced and environment averaged Raman spectra and transcriptomes, we conducted a leave-one-out cross-validation. One measurement from environment *ε_i_* was removed, and PLS-R was applied to conduct a linear regression against the remaining data set. Specifically, this equates to finding a matrix **A**_–*i*_ such that

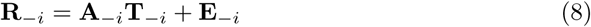

where 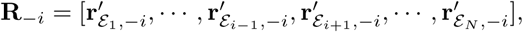, **T**_–*i*_ = [**r**′_*ε*_1__, …, **t**_ε_*i*-1__, **t**_*ε*_*i*+1__, …, **t**_ε_N__] and **E**_–*i*_ is the error matrix. Here, **r**′_*ε*, –*i*_ are the dimension reduced Raman spectra where PC-LDA against Raman spectra excluding environment *i* was applied. Also, **r**′_ε, –*i*_ and **t**_*ε*_ are mean centered by subtracting the average of the included *N* – 1 conditions. Now, when *N* – 1 < dim **t**_*ε*_ ordinary least squares regression cannot be connented to find **A**_−*i*_(In our story, for *S. pombe*, *N* – 1 = 9 < dim **t**_*ε*_ = 6560 and for *E. coli,* N – 1 = 4 < dim **t**_*ε*_ = 4349). Therefore, we applied PLS-R, which reduces the dimension of **t**_*ε*_ to below *N* – 1 so that a linear regression can be conducted, while retaining the linearity between **r**′_*ε*,-*i*_ and **t**_*ε*_ [15–17]. For all PLS-R analyses in this study, the dimensions were reduced to *N* – 3. Consequently, in our attempt in Fig. 2F to find the required numbers of transcripts to observe a linear correspondence, the number of included transcripts was increased from *N* – 3 = 10 – 3 = 7.

Once **A**_–*i*_ is estimated, **r**′_*ε*,-*i*_ can be estimated as 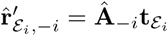 However, 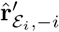 is predicted on to the PC-LDA space where environment *ε_i_* is excluded. Therefore, this space is not designed to evaluate Raman spectra obtained from environment *ε_i_*. Thus, to evaluate the estimated spectra, the basis of 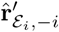 was changed to the PC-LDA space including environment *ε_i_* by 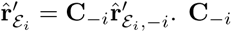 was calculated as 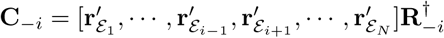, where 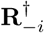 is the Moore-Penrose pseudoinverse of **R**_–*i*_. This was repeated for all *i* = 1,…,N, and PRESS_**r**_ was calculated.

To predict the transcriptome of environment *ε_i_*, we obtained the Moore-Penrose pseudoinverse of Â_–*i*_ denoted as 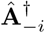, and predicted it as 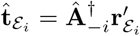. Note that in general, the estimate of transcriptome 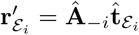 that satisfies 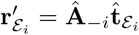 is

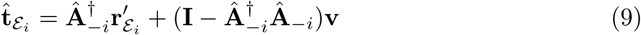

where **v** is an arbitrary vector, meaning that 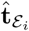 in principle cannot be determined uniquely from **r**′_*ε_i_*_. However, 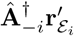 is the term that can be determined experimentally. Also, the terms 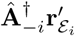 and 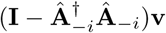 are orthogonal 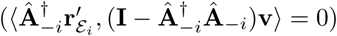, meaning that removing the term 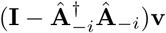 does not affect the subspace spanned by 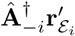. Therefore, we estimated 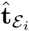 by 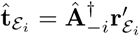. Again, this was repeated for all *i* = 1,…,*N*, and PRESS_**t**_ was calculated.

VIP scores in Table 1 and 2, and explained variances by the number of dimensions used in Fig 3A were calculated as the average of *i* = 1,…, *N* when conducting the above leave-one-out cross-validation.

### Evaluating the significance of Raman-transcriptome linearity by the permutation test

To test the significance of the Raman-transcriptome linearity, we conducted the permutation test [18, 19]. In short, false data sets were created by randomly permuting environmental assignments of transcriptomes, PRESS_**r**_ or PRESS_**t**_ values were calculated, and *p*-values of accidentally obtaining PRESS values equal to or lower than the original value were obtained.

When the number of environment conditions *N* is larger than 8, the number of possible random permutations exceeds 8! = 40, 320, and becomes computationally intensive to calculate *p*-values. Therefore, when *N* > 8, unless otherwise stated, *p*-values were calculated by randomly generating 10,000 permutations, and when *N*<8, all possible permutations were generated, *p*-values calculated from randomly generated permutations are known to underestimate exact *p*-values [20]. Therefore, when *N* > 8, *p*-values were calculated by applying the following correction: (*b* + 1)/(*m* + 1) where *b* is the number of permutations that gave PRESS values equal to or lower than the original PRESS value, and *m* is the total number of permutations [20].

## Acknowledgments

We thank the National Institute for Materials Science, Molecule & Material Synthesis Platform, for sharing the Raman microscope facilities during the initial stage of this study; Joseph Kirschvink for reading the manuscript and valuable comments; Edo Kussell and members of the Wakamoto lab for in-depth discussions. This work was supported by the Platform for Dynamic Approaches to Living System from Ministry of Education, Culture, Sports, Science and Technology Japan and Japan Agency for Medical Research and Development (to Y.W. and K.O.); Japan Society for the Promotion of Science KAKENHI (grant number 15KT0075, 15H05746 to Y.W.); and Cooperative Research Grant of the Genome Research for BioResource, NODAI Genome Research Center, Tokyo University of Agriculture (to Y.K., S.Y.). K.J.K.-K. and K.N. was supported by Grant-in-Aid for Japan Society for the Promotion of Science Fellows (grant number 17J08992 and 17J07408).

## Author contributions

K.J.K.-K. and Y.W. conceived the work. K.J.K.-K. performed Raman measurements. K.J.K.-K. and H.N. designed the culture conditions and prepared the cell samples for *S. pombe* experiments. K.N., H.F. and H.M. designed the culture conditions and prepared the cell samples for *E. coli* experiments. K.J.K.-K. and A.O. obtained RNA-Seq transcriptome data for *S. pombe*. Y.K. and S.Y. obtained RNA-Seq transcriptome data for *E. coli*. K.J.K.-K. and K.F.K. analyzed data. K.J.K.-K., H.N., A.O., K.F.K., H.M., K.O. and Y.W. evaluated the results and provided the interpretation. K.J.K.-K., K.F.K. and Y.W. wrote the manuscript. All the authors read, commented on, and approved the manuscript.

